# The floral homeotic protein SEPALLATA3 recognizes target DNA sequences by shape readout involving a conserved arginine residue in the MADS-domain

**DOI:** 10.1101/133678

**Authors:** Sandra Gusewski, Rainer Melzer, Florian Rüempler, Christian Gafert, Güenter Theiβen

## Abstract

SEPALLATA3 of *Arabidopsis thaliana* is a MADS-domain transcription factor and a central player in flower development. MADS-domain proteins bind as dimers to AT-rich sequences termed ‘CArG-boxes’ which share the consensus 5’-CC(A/T)_6_GG-3’. Since only a fraction of the abundant CArG-boxes in the *Arabidopsis* genome are bound by SEPALLATA3, more elaborate principles have to be discovered to better understand which features turn CArG-box sequences into genuine recognition sites. Here, we investigated to which extent the shape of the DNA contributes to the DNA-binding specificity of SEPALLATA3. We determined *in vitro* binding affinities of SEPALLATA3 to a variety of DNA probes which all contain the CArG-box motif, but differ in their DNA shape characteristics. We found that binding affinity correlates well with certain DNA shape features associated with ‘A-tracts’. Analysis of SEPALLATA3 proteins with single amino acid substitutions in the DNA-binding MADS-domain further revealed that a highly conserved arginine residue, which is expected to contact the DNA minor groove, contributes significantly to the shape readout. Our studies show that the specific recognition of *cis*-regulatory elements by plant MADS-domain transcription factors heavily depend on shape readout mechanisms and that the absence of a critical arginine residue in the MADS-domain impairs binding specificity.

## INTRODUCTION

The specific molecular interactions between DNA and transcription factors (TFs) are of vital importance to control gene expression. However, the determinants of target gene specificity are still poorly understood. One prominent role can certainly be attributed to the sequence of the DNA binding site, i.e. the preference for a specific nucleotide at a specific position. This mechanism of conferring protein-DNA interaction specificity is termed base readout (Rohs et al., 2010). In this case, amino acid side chains of the TFs interact via hydrogen bonds or hydrophobic interactions with bases or base pairs on the DNA and ‘read’ the sequence of the binding site (Rohs et al., 2010).

However, it was suggested that beyond the nucleotide sequence, the three-dimensional structure of the DNA may play a so far underappreciated role in governing DNA-binding specificity (Rohs et al., 2009; Parker and Tullius, 2011; Abe et al., 2015). Many transcription factors may thus not only employ a ‘base readout’ mechanism in which the primary sequence of the bases on the DNA is recognized, but also a ‘shape readout’ of the DNA to achieve binding specificity (Rohs et al., 2010).

Shape readout (sometimes also termed indirect readout) describes the recognition of the sequence-dependent conformation and deformability of the DNA by DNA-binding proteins (Olson et al., 1998; Rohs et al., 2010; Watkins et al., 2010). One shape readout mechanism that has recently received particular attention is the recognition of the width of the minor groove (Rohs et al., 2009). In contrast to the major groove, the minor groove of the DNA possesses relatively few base-pair specific opportunities for hydrogen bonding (Seeman et al., 1976; Rohs et al., 2010). However, depending on the DNA sequence the width of the minor groove can substantially vary. It was suggested that these structural differences contribute to the DNA-binding specificity of many transcription factor families (Reeves and Beckerbauer, 2001; Joshi et al., 2007; Rohs et al., 2009; Slattery et al., 2011).

Most TFs probably use an interplay between shape and base readout mechanisms to recognize DNA-binding sites but it is largely unclear to which extent shape vs. base readout contribute to protein-DNA interactions for different transcription factor families (Slattery et al., 2014). The different modes of protein-DNA recognition do however have a profound influence on the interpretation and prediction of binding site preferences. For TFs that employ primarily a base readout mechanism, individual nucleotides of the DNA-binding site contribute to the overall binding affinity in a largely additive manner (Stormo and Zhao, 2010; Zhao and Stormo, 2011). Binding motifs can in this case be accurately depicted by consensus sequences or position weight matrices. In contrast, shape readout requires a specific sequence-dependent 3D DNA structure which can only be realized by certain DNA sequences (Parker and Tullius, 2011). Accordingly, shape readout is often connected with the occurrence of intra-motif-dependencies which originate from physical interactions between base pairs (Zhou et al., 2015; Mathelier et al., 2016). Such dependencies are usually not accounted for in consensus binding sequences or position weight matrices which implicitly assume independence among the base pair positions of a DNA binding site. Thus, identifying whether TFs employ a shape readout mechanism can significantly improve our understanding of binding specificities and can improve the accuracy of binding site prediction models (Bauer et al., 2010; Meysman et al., 2011; Maienschein-Cline et al., 2012; Gordan et al., 2013; Dror et al., 2014; Yang et al., 2014; Abe et al., 2015; Zhou et al., 2015).

Here, we use the MIKC-type MADS-domain protein SEPALLATA3 (SEP3) from *Arabidopsis thaliana* to explore in more detail how gene regulatory proteins employ shape readout mechanisms. MADS-domain proteins are transcription factors that are present in almost all eukaryotes. They are essential transcription factors in fungi and animals (Theißen et al., 1996) and the subgroup of MIKC-type proteins constitutes a large family in plants (Schwarz-Sommer et al., 1990; Gramzow and Theißen, 2010; Smaczniak et al., 2012a). SEP3 is a key regulator of flower development. It is involved in the determination of floral organs and acts in a largely redundant manner with the closely related proteins SEP1, SEP2 and SEP4 (Mandel and Yanofsky, 1998; Pelaz et al., 2000; Ditta et al., 2004; Kaufmann et al., 2009). Consequently, *sep1 sep2 sep3* triple mutants possess ‘flowers’ entirely composed of sepal-like organs and *sep1 sep2 sep3 sep4* quadruple mutants develop leave-like organs instead of floral organs (Ditta et al., 2004).

In general, MADS-domain proteins bind as dimers to CArG-box sequence elements with the consensus sequence 5’-CC(A/T)_6_GG-3’ or very similar sequences (Schwarz-Sommer et al., 1990; Pellegrini et al., 1995; Shore and Sharrocks, 1995; Folter and Angenent, 2006; Melzer et al., 2006). X-ray crystal and NMR structures of the human MADS-domain transcription factors SRF and MEF2A revealed that DNA contacts are made with the minor as well as with the major groove of the DNA (Pellegrini et al., 1995; Huang et al., 2000; Santelli and Richmond, 2000). Amino acid residues of the α-helix of the MADS-domain insert into the major groove and make base-specific contacts at the edge of the 10-base-pair CArG-box sequence and beyond (Pellegrini et al., 1995; Huang et al., 2000). In addition, some contacts of the α-helix of the MADS-domain are also made with the minor groove of DNA (Pellegrini et al., 1995; Santelli and Richmond, 2000). A region N-terminal to this α-helix also inserts into the minor groove and some amino acid residues make contacts with the A/T bases in the center of the CArG-box (Pellegrini et al., 1995; Huang et al., 2000; Santelli and Richmond, 2000). However, the contacts between this N-terminal extension of the MADS-domain and the minor groove seem to be not completely base pair-specific since both A·T and T·A base pairs appear to be accepted at the six central CArG-box positions as observed in *in vitro* assays with several MADS-domain transcription factors (Leung and Miyamoto, 1989; Pollock and Treisman, 1990; Huang et al., 1996; Riechmann et al., 1996; Acton et al., 1997; Meierhans et al., 1997; West et al., 1998).

The presence of an AT-rich core in the center of the CArG-box is especially interesting as it permits the formation of A-tracts. A-tracts have been defined as sequences with at least four consecutive A·T base pairs without an intervening TpA step (i.e. A_n_T_m_, n+m ≥ 4) (Hud and Plavec, 2003; Stefl et al., 2004). In general, A-tracts have been described to be an important component of recognition sites for eukaryotic transcription factors and prokaryotic transcriptional regulators which employ a shape readout mechanism (Rohs et al., 2009; Rohs et al., 2010). These sequences possess a special structure which is formed cooperatively (Haran and Mohanty, 2009). Specifically, A-tract sequences exhibit a very narrow minor groove. Recently, the occurrence of A-tracts has been implied to be positively correlated with binding events in SEP3 ChIP-seq experiments (Muiño et al., 2014).

Here, we present qualitative and quantitative protein binding data which support the hypothesis of shape readout by the MADS-domain transcription factor SEP3. We demonstrate that binding affinities of SEP3 to ‘perfect’ CArG-box sequences, i.e. CArG-boxes which are in agreement with the 5’-CC(A/T)_6_GG-3’ consensus, can vary substantially, depending on the presence or absence of A-tract sequences within the CArG-box. We further demonstrate that a highly conserved arginine residue in the MADS-domain of SEP3 is critical for conferring binding affinity and specificity. Our data suggest that SEP3 employs a mixture of shape and base readout mechanisms to achieve target gene specificity.

## RESULTS

### SEP3_MI_ binds with especially high affinity to CArG-boxes with A-tracts in the AT-rich core

To obtain a detailed and quantitative picture of the DNA-binding affinities of SEP3, we purified an N-terminal fragment of the protein, which contains the DNA-binding MADS-domain and the I-domain that mediates dimerization (Materials and methods and Supplemental Figures 1-4). This truncated protein, which contains the first 90 amino acids of the 251-amino-acid long native SEP3 protein, is termed SEP3_MI_ henceforth. The MADS-and the I-domain have previously been shown to be necessary and sufficient for DNA-binding of plant MADS-domain proteins (Huang et al., 1996).

**Figure 1.**
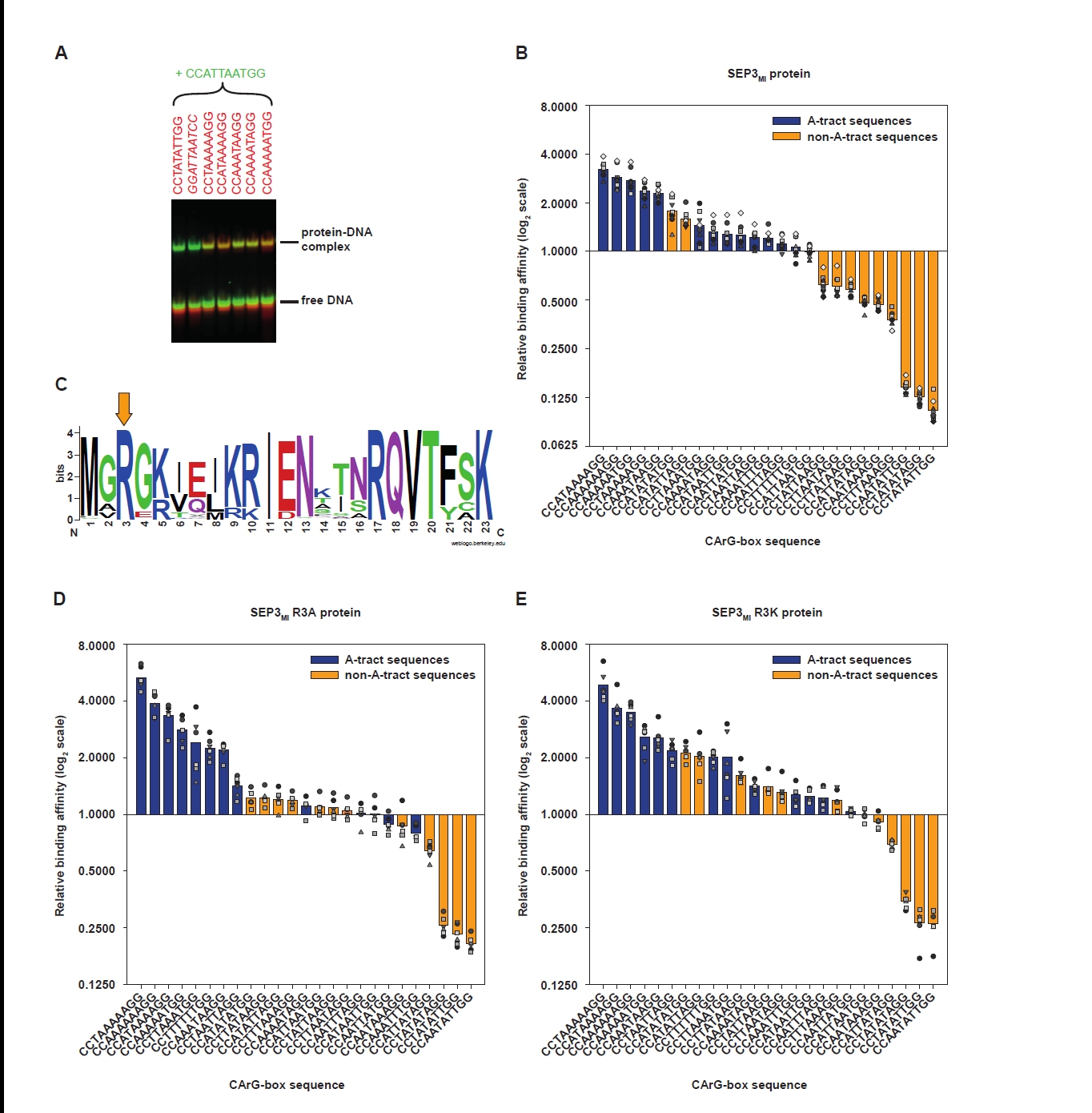
SEP3_MI_ binds preferentially to CArG-box sequences with an A-tract. **(A)** Exemplary QuMFRA assay result. Probes were incubated with purified SEP3_MI_ protein. Probes which had a Cy5-label are shown in red in the gel image. The central sequence of the probes is shown in red above the gel. Additionally, a reference probe with the central CArG-box sequence 5’-CCATTAATGG-3’ was added to each binding reaction (labelled with 6-Fam and shown in green in the gel image). **(B)** Relative affinities obtained by the QuMFRA assay are ordered from high (left) to low affinity (right) and plotted in logarithmic scale. The reference sequence used here (5’-CCATTAATGG-3’) is defined to have a relative affinity of 1. Replicates are shown by small circles, triangles, squares or diamonds, respectively. CArG-box sequences with an A-tract (blue bars) are contrasted against sequences without A-tracts (orange bars). A-tracts are defined as being sequences with a minimum of four consecutive A·T base pairs without an intervening TpA step. **(C)** Arginine R3 is highly conserved among plant MIKC-type MADS-domain proteins. The sequence logo depicts the first 23 amino acids of the MADS-domain making use of 1325 sequences of seed plant MIKC-type MADS-domain transcription factors. Arginine R3 is highlighted by the orange arrow above the sequence logo. **(D)**, **(E)** Binding specificity is impaired in the single mutants SEP3_MI_ R3A and SEP3_MI_ R3K. QuMFRA results for **(D)** SEP3_MI_ R3A and **(E)** SEP3_MI_ R3K are plotted.

To assess whether the width of the minor groove correlated with the DNA-binding of SEP3_MI_, 25 DNA probes which differ in predicted minor groove width, were designed (Supplemental Data Set 1). The DNA probes used for this assay all perfectly matched the 5’-CC(A/T)_6_GG-3’ consensus and were identical to each other outside the consensus sequence. The only difference between the probes was the number and order of adenine and thymine bases in the AT-rich core of the CArG-box. To create probes with variations in minor groove widths, we took advantage of the fact that A-tracts exhibit narrow minor grooves (Haran and Mohanty, 2009). In contrast, TpA steps tend to widen the minor groove (Haran and Mohanty, 2009). Also, in case of consecutive adenine base pairs the minor groove gets progressively narrower in 5’ to 3’ direction of the adenine strand (Haran and Mohanty, 2009). We could therefore design a number of CArG-box cores, which differ in predicted minor groove width, but otherwise have very similar sequence compositions (Table 1, Supplemental Data Set 1).

**Table 1.**
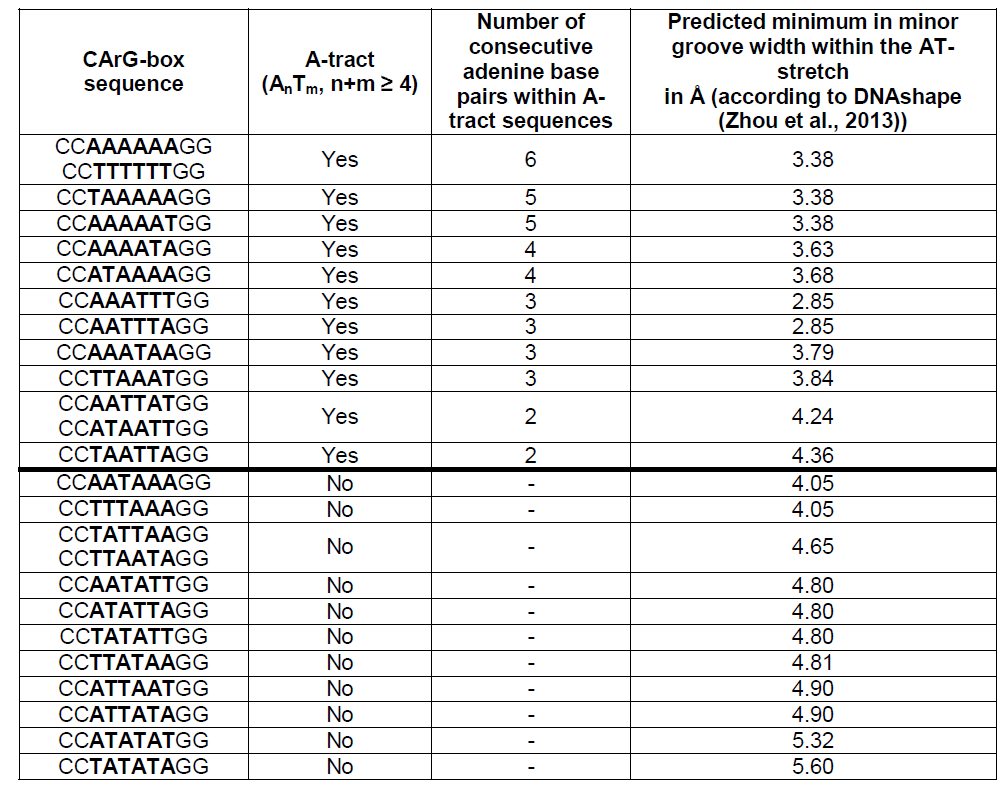
Sequences of DNA probes for the determination of SEP3_MI_ binding specificity

DNA-binding of SEP3_MI_ to these DNA probes was determined by Quantitative Multiple Fluorescence Relative Affinity (QuMFRA) assays (Figure 1A, Supplemental Data Set 2 and Supplemental Figure 5) (Man and Stormo, 2001). This is an electrophoretic mobility shift assay (EMSA) based technique that allows direct estimation of relative protein affinities to two DNA-probes (Man and Stormo, 2001).

**Figure 5.**
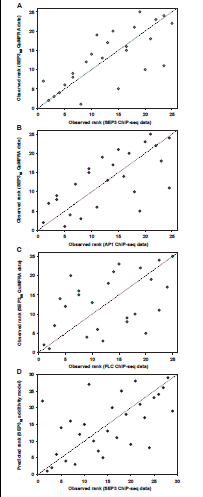
Comparison of *in vitro* QuMFRA data with *in vivo* ChIP-seq data. **(A)** Affinities of SEP3_MI_ to various CArG-box sequences, which were obtained by the QuMFRA assay, were ranked and plotted against the ranked mean ChIP-seq scores of the same CArG-box sequences for the “wild-type” (wt) SEP3 (data modified from Muiño et al., 2014). QuMFRA and ChIP-seq data show a good degree of correlation (Spearman’s rank correlation rho = 0.7462, S = 660, p-value = 1.846e-05). **(B)** Scatter plot of *in vitro* measured SEP3_MI_ binding affinities against AP1 ChIP-seq data (data modified from Muiño et al., 2014). SEP3_MI_ binding affinities and AP1 ChIP-seq data correlate with Spearman’s rank correlation rho = 0.6754, S = 844, p-value = 2.118e-04. **(C)** SEP3_MI_ binding affinities are plotted against FLC ChIP-seq data (data modified from Muiño et al., 2014). SEP3_MI_ QuMFRA data and FLC ChIP-seq data correlate with Spearman’s rank correlation rho = 0.4676, S = 1384, p-value = 1.843e-02. **(D)** Scatter plot of predicted (by the additive model) versus observed (in ChIP-seq data) binding site ranks. Values predicted by the additive model and ChIP-seq data correlate significantly (Spearman’s rank correlation rho = 0.5744, S = 1728, p-value = 1.12e-03).

We found that relative binding affinities of SEP3_MI_ to different CArG-boxes varied up to a factor of 30 (Figure 1B). The highest relative affinity was measured for 5’-CCATAAAAGG-3’ and the lowest for 5’-CCTATATTGG-3’.

The substitution of adenine or thymine against cytosine or guanine within the AT-rich core of the CArG-box decreased binding affinity of SEP3_MI_ (Supplemental Data Set 2) as has been described previously for SRF (Leung and Miyamoto, 1989). However, mutations within the AT-stretch differed in the magnitude of their negative effect. The relative binding affinity of SEP3_MI_ to the CArG-box sequence 5’- CCAAGAAAGG-3’ was ten times higher than the affinity of 5’-CCAACAAAGG-3’. The affinity to 5’-CCAAGAAAGG-3’ was even higher than to the ‘perfect’ CArG-boxes 5’-CCTATATAGG-3’ or 5’-CCAATATTGG-3’ (Supplemental Data Set 2). As negative control, a mutated sequence where the ‘CC’ and ‘GG’ borders of the CArG-box were reversed (5’-GGATTAATCC-3’) was used. The affinity to this probe was so low that it could hardly be detected by the QuMFRA assay (Supplemental Data Set 2).

13 of the 25 probes that perfectly matched the 5’-CC(AT)_6_GG-3’ consensus contained an A-tract, whereas 12 sequences did not. Intriguingly, 10 out of 12 non-A-tract sequences showed a lower affinity for binding to SEP3_MI_ than the A-tract containing sequences (Figure 1B). In general, the binding affinity of A-tract sequences versus non-A-tract sequences is significantly different (Wilcoxon rank sum test, W = 140, p-value = 3.438e-04). This preference of SEP3_MI_ for A-tract sequences in the CArG-box core is in agreement with the hypothesis that SEP3 recognizes the structure of the DNA via a shape readout mechanism and that the protein preferentially binds to CArG-box sequences with a narrow minor groove. These data are also in agreement with the hypothesis that position interdependence among nucleotides of the DNA-binding site exists.

To analyse whether positions flanking the CArG-box influence the preference of SEP3_MI_ for A-tracts within the CArG-box, we studied relative binding affinities to probes that differed in 6 nucleotides on each side immediately adjacent to the CArG-box. Of the three different flanking sequences tested, all showed similar relative affinities for sequences with different CArG-box cores, including a preference for A-tract containing CArG-box cores (Supplemental Data Set 2, Supplemental Figure 6). Importantly, however, the overall affinity was severely affected by the flanking sequences with relative affinities varying by ca. 50fold between different flanking sequences (Supplemental Data Set 2), indicating that a sequence motif longer than the CArG-box is important for high affinity binding of SEP3.

### SEP3_MI_ binds with high affinities to CArG-boxes

We also tested DNA-binding of SEP3_MI_ using saturation-binding assays in which a constant amount of DNA probe was titrated against increasing amounts of SEP3_MI_. Two CArG-boxes, both in perfect agreement with the 5’-CC(A/T)_6_GG-3’ consensus sequence (i.e. 5’-CCAAAAAAGG-3’ and 5’-CCATTAATGG-3’) were tested and compared (Supplemental Figure 7). The apparent association constant for 5’-CCAAAAAAGG-3’ was 70 × 10^6^ M^-1^ (i.e. the dissociation constant was 15 nM), approximately twice as high as for 5’-CCATTAATGG-3’ (Table 2). This was close to the binding difference detected by QuMFRA. Our measured apparent association constants were in the same range as the published affinity constant for full-length SEP3 binding to a single CArG-box (5’-CCAAATAAGG-3’) which was determined as 150 × 10^6^ M^-1^ (Jetha et al., 2014). We also tested binding to the mutated sequence in which the ‘CC’ and ‘GG’ borders of the CArG-box were reversed. As expected, binding to this sequence was very weak and therefore determination of the association constant was not possible (Supplemental Figure 7B). Overall, these results confirm the binding differences detected in the QuMFRA assay. They also show that SEP3_MI_ binds with high affinity to DNA, as would be expected for a transcription factor. Dissociation constants of transcription factors typically range from 1·10^−9^ M (nM) to 1·10^−13^ M (0.1 pM) (Schleif, 2013).

**Table 2.** Affinity and dissociation constants of SEP3_MI_ dimers binding to a CArG-box. For each protein and probe the arithmetical means of four replicates are given with the respective standard deviation.

| Protein | DNA probe sequence |  |  |  |
| --- | --- | --- | --- | --- |
|  | 5'-CCAAAAAAGG-3' |  | 5'-CCATTAATGG-3' |  |
|  | K<sub>A,dim</sub> (x 10<sup>6</sup> M<sup>-1</sup>) | K<sub>D,dim</sub> (nM) | K<sub>A,dim</sub> (x 10<sup>6</sup> M<sup>-1</sup>) | K<sub>D,dim</sub> (nM) |
| SEP3<sub>MI</sub> | 70 ± 15 | 15 ± 3 | 30 ± 7 | 35 ± 10 |
| SEP3<sub>MI</sub> R3A | 10 ± 5 | 113 ± 42 | 2.3 ± 1.1 | 500 ± 213 |
| SEP3<sub>MI</sub> R3K | 20 ± 8 | 56 ± 20 | 2.2 ± 0.5 | 463 ± 92 |

### A conserved arginine in the MADS-domain confers binding specificity

It is known that especially arginine residues in DNA-binding domains are involved in the shape readout of the DNA by binding to exceptionally narrow minor grooves (Rohs et al., 2009). In the structurally characterized MADS-domains of SRF and MEF2A, an N-terminal arm extends deep into the minor groove of the DNA (Pellegrini et al., 1995; Huang et al., 2000; Santelli and Richmond, 2000). The N-terminal arm possesses a highly conserved arginine residue at the third position of the MADS domain (R3) that is involved in DNA-binding in SRF and MEF2A (Pellegrini et al., 1995; Huang et al., 2000; Santelli and Richmond, 2000) and that is also present in almost every plant MIKC-type MADS-domain protein including SEP3 (Figure 1C).

In order to elucidate the role of the arginine residue R3 in binding affinity and specificity, we substituted it in SEP3_MI_ by either a lysine (which is chemically very similar to arginine) or an alanine (which lacks the amino acid side chain beyond the β carbon) residue, creating the proteins SEP3_MI_ R3K or SEP3_MI_ R3A, respectively.

Both mutant proteins showed a reduced DNA-binding affinity in comparison to the wild-type SEP3_MI_ protein in a saturation binding assay (Table 2 and Supplemental Figure 7). However, there was no strong difference in binding affinity between the lysine- and the alanine-substituted protein for the two CArG-box sequences tested. These results indicate that the arginine residue is critical for obtaining high DNA-binding affinities.

To test binding specificity of the mutated proteins, both variants were tested with the QuMFRA assay (Figure 1D and 1E). SEP3_MI_ R3A and SEP3_MI_ R3K still significantly prefer CArG-box sequences containing an A-tract over non-A-tract sequences (Wilcoxon rank sum test, W = 123, p-value = 1.352e-02 (for SEP3_MI_ R3A) and W = 128, p-value = 5.48e-03 (for SEP3_MI_ R3K)). The sequences with the highest affinities contained an A-tract, whereas the probes with the lowest affinities did not contain an A-tract. However, among the CArG-box sequences with moderate affinities there are both A-tract and non-A-tract sequences. In the case of SEP3_MI_ R3A there are many probes which hardly differ in their affinity (Figure 1D). For SEP3_MI_ R3K binding specificity is altered in a way that several non-A-tract sequences have a higher affinity than sequences with an A-tract (Figure 1E).

In contrast to the wild-type SEP3_MI_ protein the mutant proteins SEP3_MI_ R3A and SEP3_MI_ R3K are less able to differentiate between A-tract and non-A-tract sequences. For the wild-type SEP3_MI_ the average affinity constant for A-tract sequences was 2.74 times higher than for non-A-tract sequences, however, for the mutant proteins the ratio decreased to a value of 2.55 for SEP3_MI_ R3A and 2.07 for SEP3_MI_ R3K, respectively.

### Insertion of modified bases supports the shape readout mechanism in the minor groove

To further investigate into the importance of the shape readout of the DNA in the minor groove by SEP3_MI_, we analyzed binding of the protein to DNA probes containing the non-standard bases hypoxanthine (abbreviated with “I” because hypoxanthine is the nucleobase of the nucleoside inosine) or diaminopurine (D). Hypoxanthine can pair with cytosine, while diaminopurine base pairs with thymine. Intriguingly, the pattern of hydrogen bond donors and acceptors presented by an I·C pair is identical with that of a G·C base pair in the major groove and to that of an A·T base pair in the minor groove (Supplemental Figure 8) (Wang et al., 1998). In contrast, the pattern of hydrogen bond donors and acceptors presented by a D·T base pair is identical with that of an A·T base pair in the major groove and to that of a G·C base pair in the minor groove (Supplemental Figure 8) (Bailly et al., 1995).

We found that the substitution of an A·T base pair by a G·C base pair within the AT-stretch of the CArG-box heavily decreased the binding affinity of SEP3_MI_ (Figure 2A, B). However, when an I·C base pair was introduced binding affinity could almost completely be restored (Figure 2A, B). In contrast, the binding affinity of a probe with a D·T base pair was similar to the corresponding probe with a G·C base pair and markedly lower as compared to a probe with an A·T base pair at that position (Figure 2B). When the complete AT-stretch was replaced by a GC-stretch, binding was nearly abolished. However, when the AT-stretch was substituted by six consecutive I·C base pairs, binding could be partly restored (Figure 2C).

**Figure 2.**
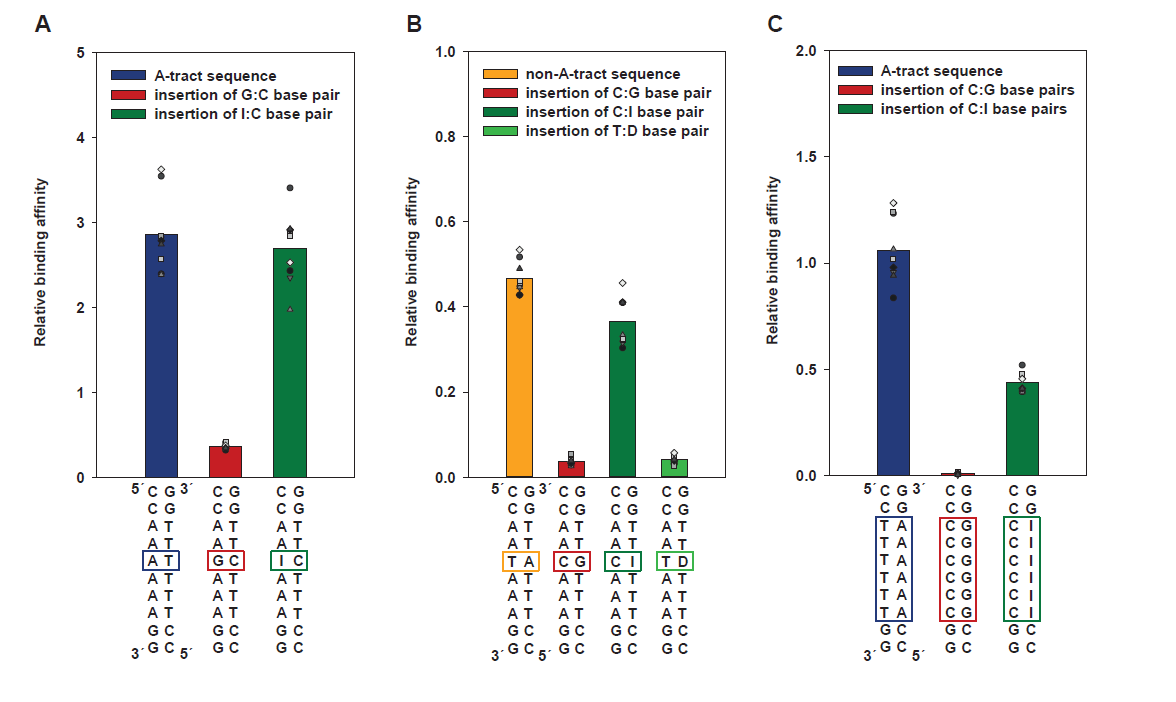
Insertion of modified bases supports the hypothesis of a shape readout mechanism in the minor groove. Relative affinities were obtained by the QuMFRA assay. 5’-CCATTAATGG-3’ was used as the reference sequence. CArG-box sequences with G·C base pairs (red bars) are compared with A·T base pairs (blue or orange bars, respectively), I·C base pairs (dark green bars in **(A)**, **(B)**, **(C)**), where “I” stands for the base hypoxanthine/ the nucleoside inosine, and D·T base pairs (light green bar in **(B)**), where “D” denotes the base diaminopurine. Replicates are shown by small circles, triangles, squares or diamonds, respectively.

These findings support the hypothesis that the hydrogen bond pattern of A·T base pairs in the minor groove of the CArG-box center - with its structural consequences for minor groove geometry (see also discussion below) – is very important for binding specificity of SEP3_MI_.

### A simple model of additivity describes binding specificities of SEP3_MI_ R3A and SEP3_MI_ R3K better than that of SEP3_MI_

Based on our affinity measurements we aimed at developing a model which can explain SEP3_MI_ binding specificity. The simplest assumption would be that each base pair of the binding site contributes independently to the overall binding energy, irrespective of the identity of the neighboring positions (Stormo and Fields, 1998; Stormo and Zhao, 2010).

In this framework, experimentally determined affinities of a reference probe and of probes that deviate from this reference in a single base pair allow to calculate the binding energy contribution of each base pair. This can subsequently be used to predict binding energies for probes that deviate in more than one base pair from the reference probe, as previously described (Stormo and Fields, 1998; Maerkl and Quake, 2007).

When we used such a model and applied it to our affinity measurements for SEP3_MI_ (Supplemental Data Set 3), predicted binding energies correlated with experimentally observed binding energies with a Spearman’s rank correlation coefficient rho of 0.73 (Figure 3A). The root mean square error of prediction (RMSEP) has a value of 1.36 kJ/mol.

**Figure 3.**
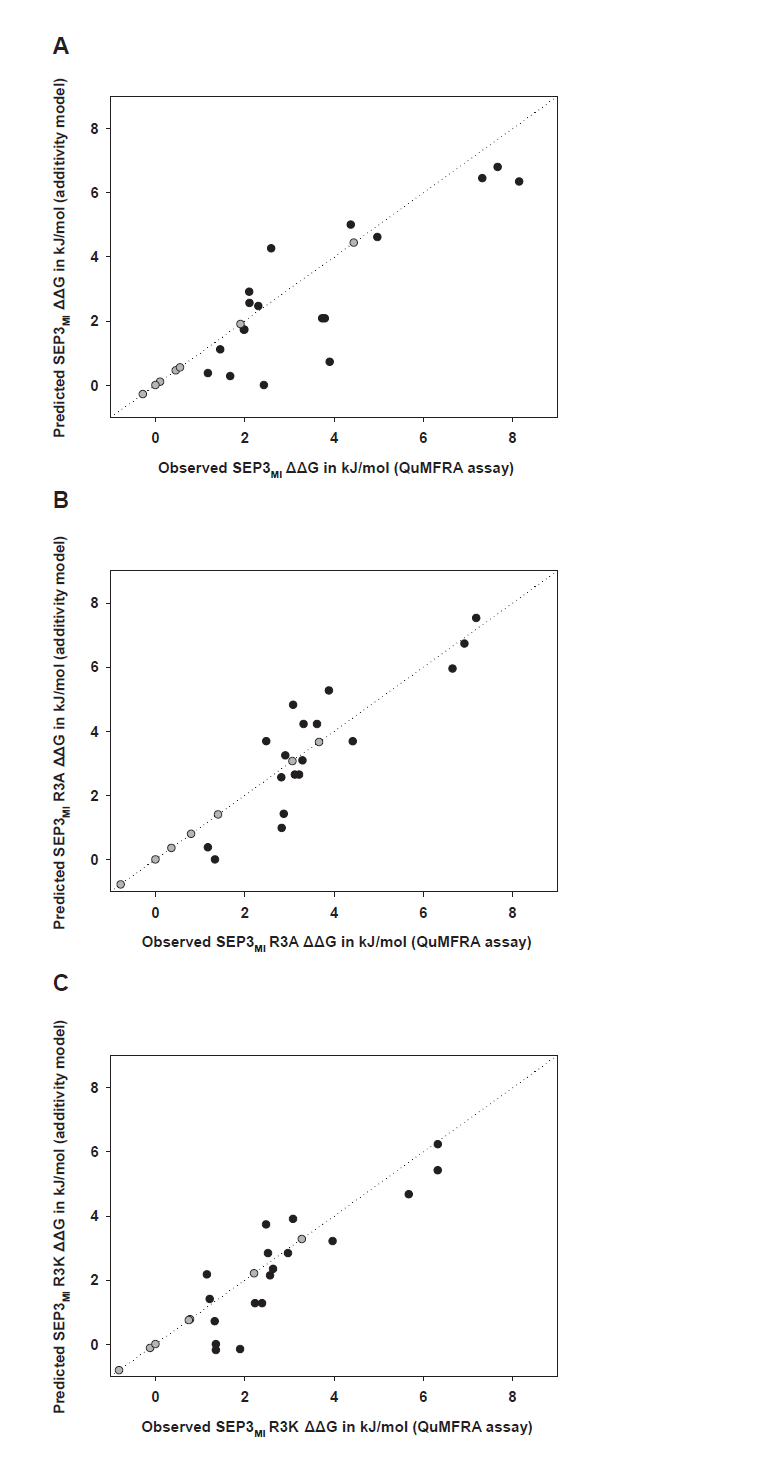
Additivity model. The dotted line marks the expected 1:1 ratio for predicted and observed values. Changes in the Gibbs free energy (ΔΔG) observed in the QuMFRA assay and predicted values based on the additive model are compared. The reference sequence 5’-CCAAAAAAGG-3’ as well as the values for sequences with single A-to-T substitutions are plotted (gray dots), on which the predictions for the other CArG-box sequences (black dots) are based. **(A)** Additivity model for SEP3_MI_. There is a significant correlation (Spearman’s rank correlation rho = 0.7324, S = 259, p-value = 5.47e-04). **(B)** Additivity model for SEP3_MI_ R3A. There is a significant correlation (Spearman’s rank correlation rho = 0.8254, S = 169, p-value = 2.47e-05). **(C)** Additivity model for SEP3_MI_ R3K. There is a significant correlation (Spearman’s rank correlation rho = 0.8151, S = 179, p-value = 3.779e-05).

Interestingly, for the mutant proteins SEP3_MI_ R3A and SEP3_MI_ R3K higher Spearman’s rank correlation coefficients were obtained for the comparison between the QuMFRA measurements and the predictions from the additive model with a value of 0.83 for SEP3_MI_ R3A and 0.82 for SEP3_MI_ R3K (Figure 3B and 3C). In addition, the RMSEP values for the additive models of SEP3_MI_ R3A and of SEP3_MI_ R3K are lower than for SEP3_MI_ (0.99 kJ/mol for SEP3_MI_ R3A and 0.97 kJ/mol for SEP3_MI_ R3K).

Taken together, this indicates that the assumption of independent contributions of the base pairs to the binding energy does reflect the protein-DNA recognition mode of SEP3_MI_ R3A and SEP3_MI_ R3K to a greater extent than that of SEP3_MI_.

### Binding affinity correlates with several DNA shape parameters

Since SEP3 prefers A-tract sequences and A-tracts are known to have characteristic structural features (Haran and Mohanty, 2009), we tried to better understand how the measured binding affinities and the structural determinants of the studied CArG-box DNA sequences (namely the minor groove width - MGW, the propeller twist - ProT, the roll angle – Roll and the helical twist - HelT) are correlated with each other. The online tool DNAshape (Zhou et al., 2013) was used to predict these structural features for every base pair or base pair step, respectively, of the DNA probes, for which we had determined SEP3_MI_ binding affinities using the QuMFRA assay (Supplemental Data Set 4).

To compare the propensity for a shape readout mechanism of SEP3_MI_ with that of the mutant proteins SEP3_MI_ R3A and SEP3_MI_ R3K, we tested how the 25 measured binding affinities of the respective proteins to the different CArG-box sequences correlated with a single value per CArG-box sequence and per DNA structural parameter. For those analyses the minimal value of minor groove width, propeller twist and roll angle and the maximal value of helical twist within the AT-stretch for each CArG-box sequence were chosen (Table 3, Figure 4).

**Table 3.** Testing for correlation between binding affinities and DNA structural parameters. Spearman’s rank correlation coefficient rho is given for the comparisons between binding affinities determined by the QuMFRA assay and the minimal or maximal value of the respective DNA structural parameter for the same CArG-box sequence. The significance level is Bonferroni-corrected for a critical value of p ≤ 0.05 (p ≤ 0.0125 after Bonferroni correction).

| DNA structural parameter | SEP3<sub>MI</sub> protein |  |  | SEP3<sub>MI</sub> R3A protein |  |  | SEP3<sub>MI</sub> R3K protein |  |  |
| --- | --- | --- | --- | --- | --- | --- | --- | --- | --- |
|  | rho | p-value | significance level | rho | p-value | significance level | rho | p-value | significance level |
| Minimum in minor groove width (MGW) | -0.5042 | 0.0102 | significant | -0.4389 | 0.0282 | not significant | -0.4656 | 0.0190 | not significant |
| Minimum in propeller twist (ProT) | -0.5748 | 0.0027 | significant | -0.5617 | 0.0035 | significant | -0.5458 | 0.0048 | significant |
| Minimum in roll angle (Roll) | -0.5682 | 0.0030 | significant | -0.4096 | 0.0420 | not significant | -0.4760 | 0.0162 | not significant |
| Maximum in helical twist (HelT) | 0.5697 | 0.0030 | significant | 0.5319 | 0.0062 | significant | 0.5008 | 0.0108 | significant |

**Figure 4.**
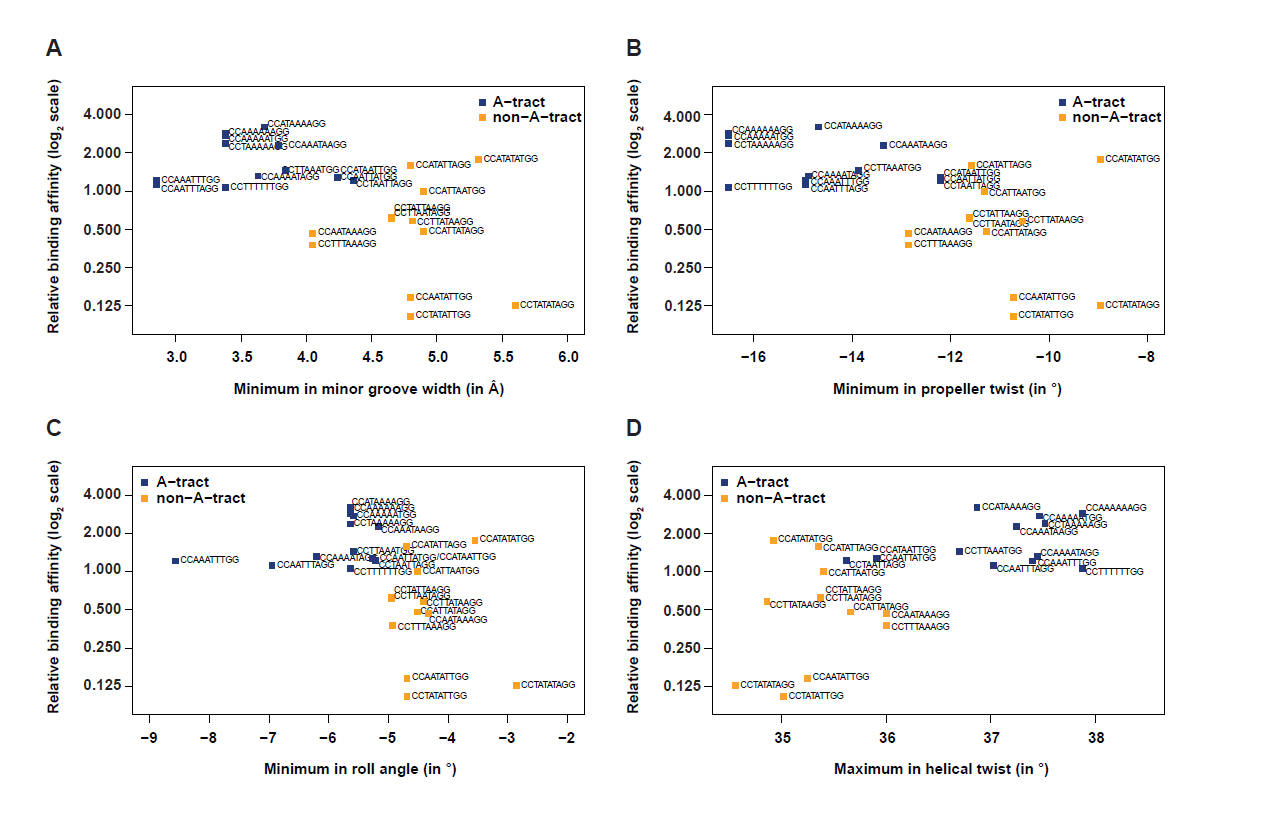
Binding affinity correlates with DNA shape parameters. Relative binding affinities of SEP3_MI_ determined by the QuMFRA assay were plotted against the minimal value of **(A)** minor groove width, **(B)** propeller twist, **(C)** roll angle and **(D)** the maximal value of helical twist of the respective DNA structural parameter for the CArG-box sequences as labeled. CArG-box sequences which contain an A-tract are labeled with squares in dark blue, non-A-tract sequences are labeled with orange squares.

Cohen’s standard (derived from Cohen’s d) (Cohen, 2003) was used to evaluate how the correlation coefficient determines the strength of the relationship, or the effect size, where coefficients between 0.10 and 0.29 represent a small association, coefficients between 0.30 and 0.49 represent a medium and coefficients above 0.50 represent a large association. The same is true for negative correlation coefficients, e.g. values below -0.50 equally represent large associations.

All correlation coefficients were in the range of medium or large associations. However, for all four DNA structural parameters the correlation is higher for SEP3_MI_ than for either SEP3_MI_ R3A or SEP3_MI_ R3K (Table 3). After Bonferroni-correction for multiple testing the correlations for the roll angle and for the minor groove width for the mutant proteins are not significant anymore, whereas for SEP3_MI_ all four structural parameters correlate significantly with binding affinity (Table 3). This implies that the mutant proteins have a weaker tendency to discriminate between different DNA conformations of the AT-stretch especially with respect to the shape of the minor groove.

The analyses also show that A-tract and non-A-tract sequences do not only differ in sequence but also in their shape characteristics, as expected. A-tract sequences did not only have higher relative binding affinities to SEP3_MI_, but they also have a smaller minimum in minor groove width, more negative propeller twist, more negative roll angles and higher helical twist values compared to non-A-tract sequences which leads to a strong clustering effect when comparing both groups (Figure 4). Thus, the preference of SEP3_MI_ for A-tract sequences can be explained by the preference for certain DNA shape features.

### Correlation of *in vivo* and *in vitro* binding specificities of SEP3

To better understand how our *in vitro* determined affinity measurements compare to *in vivo* binding of SEP3 in cell nuclei of *A. thaliana* flowers, we compared QuMFRA results obtained by this study to published ChIP-seq data for SEP3 (Supplemental Data Set 5) (Kaufmann et al., 2009; Muiño et al., 2014). To account for the different types of measurement, the values of each data set were ranked with high ranks representing high affinity or high ChIP-seq scores. A significant correlation was observed between the two datasets with a Spearman’s rank correlation rho of 0.7462 (Figure 5A). In addition, correlation with SEP3 ChIP-seq data was stronger than for ChIP-seq data of the closely related proteins AP1 with a Spearman’s rank correlation rho of 0.6754 or FLC with a correlation coefficient rho of 0.4676 (Kaufmann et al., 2010; Deng et al., 2011; Muiño et al., 2014) (Figure 5B, C, Supplemental Data Set 5).

We also compared the results of the additivity model with ChIP-seq data (Muiño et al., 2014). We evaluated how the predictions of the model and ChIP-seq score means correlated for the 29 CArG-box sequence pairs which had not been involved in training the additivity model (Supplemental Data Set 5). A statistically significant correlation was detected (Figure 5D), indicating that the additivity model is partially able to explain and predict SEP3 *in vivo* binding specificity.

## DISCUSSION

### SEP3 binds different CArG-boxes with very different affinities

We found an up to 30fold difference in the affinity (as estimated by K_A_ values) between different CArG-boxes adhering the consensus 5’-CC(A/T)_6_GG-3’ and varying only in the number and order of As and Ts in the AT-rich core of the CArG-box. It has previously been reported that protein affinity differences as little as 6fold can have significant impacts on gene regulation (Joshi et al., 2007). Thus, a 30fold difference between affinities for different CArG-boxes may have profound consequences for target gene recognition. Previous reports indicated that 18 % of the *Arabidopsis* genes contain a perfect CArG-box motif in a region 3000 bp upstream of the transcription start, a number that is probably vastly exaggerating the number of true target genes of MADS-domain transcription factors (Folter and Angenent, 2006). Our results indicate that this discrepancy arises at least partially because the 5’-CC(A/T)_6_GG-3’ consensus is a suboptimal representation for binding preferences of SEP3 and probably also for other MIKC-type proteins.

Quantitative analysis of SEP3_MI_ binding revealed a strong preference for CArG-boxes containing A-tracts in the AT-rich core. For example, the CArG-box 5’- CCATAATTGG-3’ that contains an A-tract (i.e. A_n_T_m_, n+m ≥ 4, underlined) was bound with 8fold higher affinity than the very similar CArG-box 5’-CCAATATTGG-3’ that lacks an A-tract. This *in vitro* determined binding preference of SEP3_MI_ is also in agreement with SEP3 *in vivo* ChIP-seq data indicating that SEP3 possesses an increased affinity to A-tract containing CArG-boxes with narrow minor grooves (Muiño et al., 2014).

A-tracts do naturally result in interdependencies between different bases at the binding site and such interdependencies are notoriously difficult to represent in position weight matrices and consensus sequences. This A-tract preference may thus at least partially explain the differences observed in affinity between different CArG-boxes.

Contrary to our expectations, there were two CArG-box sequences without an A-tract (5’-CCATATATGG-3’ and 5’-CCATATTAGG-3’) which had a higher relative affinity in the QuMFRA assay than most CArG-box sequences with an A-tract (Figure 1B). These two sequences may adopt a conformation which is similar to the A-tract conformation upon protein binding. It has been described that alternating AT sequences have a flexible structure. For example, complex formation of alternating AT DNA with netropsin, a minor groove binding agent, can induce a conformational change in the DNA including narrowing of the minor groove and a change in bending (Rettig et al., 2013). There are also other reports about the flexibility of TpA-steps and their ability to be accommodated in narrow minor grooves (Watkins et al., 2004; Tolstorukov et al., 2007; Rohs et al., 2009; Zhou et al., 2013). However, it remains unclear why the large difference between seemingly similar CArG-boxes like 5’-CCATATATGG-3’ and 5’-CCTATATAGG-3’ exists.

Beyond the preference for A-tracts in the center of the CArG-box, our results also indicate that the nucleotides immediately adjacent to the CArG-box can have drastic influence on binding affinities. This is illustrated by the fact that a more than 50fold affinity difference was observed between some flanking sequences. These results challenge the view of a ten-base-pair motif as the binding sequence for SEP3 and possibly also for other MIKC-type proteins. Earlier SELEX studies and recent ChIP-seq studies already suggested that SEP proteins have a preference for AT-rich sequences not only in the core of the CArG-box but also in the nucleotides flanking the CArG-boxes (Huang et al., 1996; Muiño et al., 2014; Pajoro et al., 2014), but the quantitative extent to which affinity depended on the flanking sequences in the present study was unexpected. Nonetheless, similar results were also reported for other unrelated transcription factors, suggesting that binding specificity might frequently be ‘hidden’ in the sequences flanking the core binding motif (Gordan et al., 2013; Jolma et al., 2013; Yang et al., 2014; Jolma et al., 2015; Nettling et al., 2015; Nitta et al., 2015) and future studies are required to shed light on the relevance of the sequences flanking the CArG-box for binding specificity.

### SEP3 employs a shape readout mechanism

A-tracts have very distinct structural features and have been repeatedly implicated to be involved in a shape readout mechanism for protein-DNA recognition (Rohs et al., 2009; West et al., 2010; Gordon et al., 2011; Hancock et al., 2013). Indeed, several DNA structural parameters typical for A-tracts correlated very well with SEP3 binding affinity (Figure 4). By substituting A·T base pairs with I·C or D·T base pairs in the CArG-box core, we could distinguish between the relative influences of the major and the minor groove of the AT-rich center on binding affinity. Introduction of I·C base pairs had only minor influences on binding affinity, whereas introduction of D·T base pairs strongly diminished binding (Figure 2). It has been described that I·C base pairs within AT-rich sequences promote minor groove compression to a similar degree as A·T base pairs would do at the same position (Hancock et al., 2013). On the other hand, D·T base pairs lead to minor groove widening very similar to G·C base pairs (Bailly et al., 1995; Bailly and Waring, 1995; Mollegaard et al., 1997). Taken together, our data indicate that SEP3 employs a shape readout mechanism for the specific binding of the AT-rich CArG-box center by detecting the special biophysical properties of the minor groove.

In several DNA-binding proteins especially arginine has been implicated in mediating shape readout by protruding deep into the minor groove (Rohs et al., 2009; Cordeiro et al., 2011; Gordon et al., 2011; Le et al., 2011). X-ray crystal structures of the human MADS-domain proteins showed that also in those cases an arginine at position 3 of the MADS-domain is involved in minor groove binding (Pellegrini et al., 1995; Huang et al., 2000; Santelli and Richmond, 2000). In order to test the biochemical consequences of a mutation of this seemingly important arginine residue, we created mutant proteins with a substitution of this arginine R3. Indeed, substitution by lysine or alanine changed binding specificity and weakened the binding preference for A-tracts. Statistical tests also indicated why the SEP3_MI_ R3A and R3K mutant proteins exhibit a change in binding specificity: their binding affinity is less dependent on the shape of the DNA (Table 3). In addition, a simple sequence-dependent additivity model captures binding specificity of the mutant proteins better than that of the wild-type protein (Figure 3). This model focuses on base readout mechanisms and does not take into account interdependencies among base pairs and hence shape readout mechanisms. Together, this suggests that SEP3_MI_ R3A and SEP3_MI_ R3K mutant proteins do employ a base readout mechanisms to a greater extent than the wild-type protein and that R3 is a critical residue for conferring shape readout. Importantly, however, also the SEP3_MI_ R3A and SEP3_MI_ R3K mutant proteins still show a preference for A-tracts within the CArG-box core and binding correlates with certain physical parameters of the DNA (Table 3), suggesting that other amino acid residues also contribute to the shape readout mechanism of SEP3.

The evolutionary conservation of the arginine residue R3 further suggests that also other MIKC-type proteins and potentially also non-MIKC-type MADS-domain proteins use shape readout for protein-DNA recognition. The involvement of an arginine residue in the shape readout of a narrow minor groove seems to be a common phenomenon for many transcription factors (Rohs et al., 2009; Rohs et al., 2010). Interestingly, this type of protein-DNA interaction occurs in prokaryotes as well as in eukaryotes and also in completely unrelated DNA-binding protein families (Rohs et al., 2009) with an increasing number of representatives being described in recent years (Mendieta et al., 2012; Quade et al., 2012; Alanazi et al., 2013; Porrua et al., 2013; Stevenson et al., 2013; Chintakayala et al., 2015).

### *In vitro* and *in vivo* binding specificity of SEP3 are strongly correlated

The comparison of *in vitro* and *in vivo* data concerning the binding specificity of SEP3 is complicated by several factors. *In vivo*, protein-protein interactions and the cooperative binding with other transcription factors may significantly alter DNA-binding specificity (Slattery et al., 2014). Especially the interaction between SEP3 and different MADS-domain transcription factors in the formation of floral quartet-like complexes may influence target gene specificity (Immink et al., 2009; Smaczniak et al., 2012b). Also, *in vivo* SEP3 may interact with multiple different partners other than MADS-domain proteins (Smaczniak et al., 2012b; Wu et al., 2012). ChIP-seq derived binding sites thus very likely represent a mixture of different SEP3-containing complexes binding to DNA (Kaufmann et al., 2009). An important issue *in vivo* is also the accessibility of the DNA binding sites to the transcription factors due to nucleosome formation (Slattery et al., 2014). In contrast, we studied the DNA binding properties of SEP3_MI_ homodimers to ‘naked’ DNA in our *in vitro* assays.

Furthermore, due to the purification strategy, our SEP3_MI_ protein lacked the K- and the C-domain and contained two additional amino acids N-terminal to the MADS-domain and we cannot exclude that this may have an influence on binding specificity. This may also cause differences between *in vitro* and *in vivo* binding.

All those differences notwithstanding, our *in vitro* data correlated very well with the published ChIP-seq data (Kaufmann et al., 2009; Muiño et al., 2014). This illustrates that intrinsic biophysical properties of the MADS-and the I-domain of SEP3 contribute significantly to *in vivo* target gene specificity of the protein. More importantly, our *in vitro* data correlated better with ChIP-seq data from SEP3 as compared to those from FLC or AP1. This suggests that intrinsic biophysical differences in DNA-binding specificity do account for differences in target gene specificity between SEP3, AP1 and FLC and hence may partially explain differences in functional specificity between different MADS-domain proteins. This is in contrast to earlier reports based on overexpression of hybrid proteins consisting of the DNA-binding part of the MADS-domain of one protein and the remaining half of the MADS-domain, the I-, K- and C-domain of another protein. Those analyses found that the functional specificity is to a large extent independent of DNA-binding specificity (Riechmann and Meyerowitz, 1997). Additional studies, e. g. using hybrid or mutant proteins under control of native MADS-domain protein promoters, may shed light on the question to which extent the DNA-binding specificity as conferred by the MADS-domain contributes to the functional specificity of the protein.

### A-tract binding of SEP3 may facilitate its role as a pioneer factor and molecular switch

There is evidence that SEP3 can act as a pioneer transcription factor (Pajoro et al., 2014), which describes the ability to bind nucleosome-associated DNA. These binding events lead to the opening of chromatin for other transcription factors either directly (by displacing nucleosomes) or indirectly (by recruiting chromatin remodeling factors) (Slattery et al., 2014). A hypothesis as to how SEP3 might displace nucleosomes is proposed by the “nucleosome mimicry model” (Theißen et al., 2016). This model states that floral quartet-like complexes (FQCs), protein complexes in which SEP3 can participate, mimic half-nucleosomes because both protein complexes, FQCs and half-nucleosomes, consist of four proteins with a similar protein complex size. In addition, DNA-binding and –wrapping of both complexes might also be similar (Theißen et al., 2016).

Interestingly, nucleosomes appear to preferentially bind to DNA sequences containing short A-tracts of ≤ 5 bp in length (Rohs et al., 2009). The fact that SEP3 prefers CArG-boxes containing A-tracts for binding suggests that histones and SEP3 may at least partially compete for the same binding sites. Indeed, at least in the human genome, promoter and enhancer regions often possess high intrinsic affinity for nucleosome assembly (Iwafuchi-Doi and Zaret, 2014). It has been suggested that this high nucleosome occupancy is a general mechanism to protect genes from accidental expression during the ‘wrong’ developmental phase and that pioneer factors with their capability to displace nucleosomes from promoters and enhancers create nucleosome free regions that are then also accessible to other transcription factors (Barozzi et al., 2014; Iwafuchi-Doi and Zaret, 2014). The abilities of SEP3 to act as both, a pioneer factor and a key developmental regulator during flower development, might therefore be intrinsically connected to each other: SEP3 may preferentially bind A-tract rich sequences to displace nucleosomes from transcription start sites; this pioneer role facilitates binding of other trans-factors that are important for gene regulation during flower development and thus establishes the function of SEP3 as a molecular switch during plant development.

## METHODS

### Protein expression and purification

*SEPALLATA3* (*SEP3*) cDNA (GenBank accession: NM_102272, positions 1-270 of the CDS) was amplified via PCR. The 270 bp fragment contains the MADS-and the I-domain and was thus termed *SEP3*_MI_ (Supplemental Figure 1A). It was cloned into the bacterial expression vector pET-15b (Merck Millipore) using NdeI and BamHI recognition sites, creating an N-terminal fused His_6_-tagged protein. The vector was then modified by deleting the CAT nucleotides of the NdeI site. *SEP3* mutants *SEP3*_MI_ R3A and *SEP3*_MI_ R3K were created by site-directed mutagenesis from the previously created vector carrying SEP3_MI_. Sequences are listed in Supplemental Table 1.

The protein expression and purification strategy was identical for all three protein variants (SEP3_MI_, SEP3_MI_ R3A and SEP3_MI_ R3K, respectively). Gel pictures and chromatograms of the protein expression and the purification steps were almost indistinguishable between the three proteins. For that reason, exemplary gel pictures and chromatograms are shown in Supplemental Figures 1-4 which are representative for all three proteins.

Proteins were expressed in Tuner DE3 *E. coli* cells, which contained the pRIL vector as a helper plasmid. Cells were grown at 37 °C in standard LB medium supplemented with 34 μg/ml chloramphenicol and 50 μg/ml carbenicillin. 4 ml of an overnight culture were used to inoculate a 50 ml preparatory culture. The preparatory culture was grown to an OD_600_ value of 0.5. 12 ml of the preparatory culture were used to inoculate the main culture with a volume of 400 ml. When the main culture reached an OD_600_ value of 0.8, protein expression was induced by the addition of IPTG to a final concentration of 0.4 mM. Afterwards, cell growth was continued at 37 °C for 30 minutes. Bacteria were collected by centrifugation for 40 min at 4 °C at a speed of 5,000 g. Pellets were stored at -80 °C. Protein expression was verified by SDS-PAGE and Coomassie brilliant blue staining (exemplary gel picture in Supplemental Figure 1B).

The bacterial pellet was thawed on ice and resuspended in 10 ml buffer containing 50 mM Tris pH 8, 300 mM NaCl, 5 mM DTT (dithiothreitol) and 10 mM imidazole. Cells were lysed by sonication on ice using a BANDELIN SONOPULS HD70 (Power: MS72/D, Cycle: 50 %). Sonication was done in eight cycles, whereby each cycle consisted of 30 seconds of sonication followed by a break of 30 seconds. The bacterial lysate was then centrifuged for 40 min at 4 °C at 20,000 g. After centrifugation the supernatant (containing soluble proteins including SEP3_MI_) was used for further purification (exemplary gel picture in Supplemental Figure 2).

Protein purification was done using an Äkta purifier 10. First, the His-tagged protein was purified on a Ni sepharose column (His-Trap FF crude, GE Healthcare) (exemplary gel picture and chromatogram in Supplemental Figure 2). The Ni sepharose column was equilibrated with a buffer containing 50 mM Tris pH 8, 300 mM NaCl, 5 mM DTT and 20 mM imidazole. After loading of the sample, the column was washed with 60 ml buffer containing 100 mM imidazole. Elution was done with 10 ml buffer with an imidazole concentration of 0.5 M and the column was finally cleaned with 9 ml buffer with an imidazole concentration of 1 M.

The eluted protein fractions were pooled and bacterial DNA was removed employing a heparin sepharose column (HiTrap Heparin HP, GE Healthcare) (exemplary gel picture and chromatogram in Supplemental Figure 3). The heparin sepharose column was equilibrated with a buffer containing 50 mM Tris pH 8, 300 mM NaCl and 5 mM DTT. After loading of the sample, a linear gradient of 15 ml from 0.300 M NaCl to 1.245 M NaCl was applied. The protein eluted at a salt concentration of approximately 0.9 M NaCl.

The eluted protein fractions were pooled and concentrated with a U-Tube concentrator (Novagen, 10 kDa molecular weight cut-off) to an approximate volume of 500 μl. Then the His-tag was removed by thrombin cleavage. For that purpose 0.6 units of thrombin (Novagen) were added. The reaction was incubated for 2 h at 25 °C and finally stopped by the addition of 10 μl 100 mM Pefabloc. The thrombin cleavage left a glycine and a serine at the N-terminus of the protein. As a result the purified SEP3_MI_ proteins have a length of 92 amino acids.

Proteins were further purified using size exclusion chromatography (Superdex75 10/300 GL column, GE Healthcare) (exemplary gel picture and chromatogram in Supplemental Figure 4). The column was equilibrated with a buffer containing 50 mM Tris pH 8, 300 mM NaCl and 5 mM DTT. SEP3_MI_ eluted after about 12-13 ml.

Again, the eluted protein fractions were pooled. The protein solution was concentrated with a U-Tube concentrator (Novagen, 10 kDa molecular weight cut-off) to an approximate volume of 100 μl.

Proteins were stored in 150 mM NaCl, 25 mM Tris pH 8, 2.5 mM DTT and 50% glycerol at either -20 °C (short-term storage) or -80 °C (long-term storage). Final protein (dimer) concentrations varied between 10 and 65 μM.

### DNA probes

Single-stranded DNA oligonucleotides for gel shift assays were ordered from biomers.net. Complementary oligonucleotides were annealed in 75 mM NaCl, 5 mM Tris-Cl (pH 8) and 0.5 mM EDTA (ethylenediaminetetraacetic acid). The probe sequences are listed in Supplemental Data Set 1.

In general, all probes were designed in the way that they contained a CArG-box of the type CC(A/T)_6_GG (bold) in the center of the probe, as for example: 5’-*AATTC*ATCGATCGTTTA**CCAAAAAAGG**AAATATCGATCG*G*-3’. The CArG-boxes are flanked by an AT-stretch on both sides (underlined), which was inferred from SELEX data for AGL2 (SEP1) (Huang et al., 1996). Nucleotides marked in italics have been designed for EcoRI cloning. The 5’-overhang (AATT) was also used for the Klenow fill-in radioactive labeling. The remaining nucleotides of the probes were chosen randomly.

Additionally, probes with a CArG-box flanked by G/C-rich stretches were tested. Further controls included probes with mutated CArG-boxes and probes with 2’-Deoxyinosine (biomers.net) and 2,6-Diaminopurine (obtained from IDT) substitutions (Supplemental Data Set 1).

For the saturation binding assay, DNA probes were radioactively labeled with [α-^32^P]-dATP in a Klenow fill-in reaction (Thermo Scientific). Labeled probes were purified using the illustra ProbeQuant kit (GE Healthcare). Labeled probes were stored at -20 °C.

For QuMFRA (Quantitative multiple fluorescence relative affinity) assays oligonucleotides labeled with a fluorescent dye (Cyanine dye 5 - Cy5, 6-Carboxyfluorescein - 6-Fam or Dyomics dye 490 - Dy-490) at the 5’-end were purchased from biomers.net, annealed with unlabeled complementary oligonucleotides and directly used in the assay.

### Electrophoretic mobility shift assays

The protein-DNA binding buffer was composed of 150 mM NaCl, 25 mM Tris-Cl (pH 8), 2.5 mM DTT, 1.5 mM EDTA, 2.5% (w/v) CHAPS (3-[(3-cholamidopropyl)dimethylammonio]-1-propanesulfonate), 0.3 mg/ml BSA (bovine serum albumin), 5% (v/v) glycerol, 1.333 mM spermidine trihydrochloride, 33.33 ng/μl Poly(deoxyinosinic-deoxycytidylic) acid sodium salt (Poly(dI-dC) • Poly(dI-dC) sodium salt) and 0.025% (w/v) Orange G. Protein (dimer) concentrations were varied between 1 nM and 3.5-5.5 μM for the saturation binding assay. For QuMFRA assays 0.1 μM to 0.4 μM protein (dimer) was used depending on the protein and the probes being tested. Concentration of the DNA probe was 0.1 nM for radioactively labeled DNA and 200 nM for probes labeled with fluorescent dyes. The volume of each binding reaction was 12 μl.

The binding reaction mixture was incubated for 2 hours at 22 °C. Samples were then loaded onto 1x TBE native 8% polyacrylamide gels which had been prepared with a 40% acrylamide/bis-acrylamide solution 19:1 and which had been pre-run for about 15 minutes. Gels were run at room temperature at 7.5 V/cm for 3 hours. For QuMFRA assays, gels were directly scanned on a FLA-7000 (Fujifilm) using the method ‘Cy5’ with laser excitation at 635 nm and filter R670 and the method ‘FAM’ (for 6-Fam and DY-490) with laser excitation at 473 nm and filter Y520. Fluorescence emission was measured with a voltage of 500 V at the photo-multiplier tube (PMT). For the saturation binding assay, gels were dried and exposed to phosphorimager screens. Afterwards screens were scanned on the imager FLA-7000 (Fujifilm). Images were quantified using the Multi Gauge Software (Fujifilm).

### Quantitative multiple fluorescence relative affinity (QuMFRA) assay

Relative binding affinities were determined using the QuMFRA assay (Man and Stormo, 2001). The apparent equilibrium association constant K_A_ of a DNA probe D is given by:

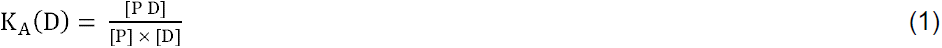

with [P D], [P] and [D] being the concentration of the protein-DNA complex, free protein and free DNA, respectively. Relative binding affinities can be obtained by calculating a ratio between the association constant K_A_ (D_1_) for the tested DNA probe D_1_ and K_A_ (D_2_) for the chosen standard DNA probe D_2_, which is part of every binding reaction in the competition assay:

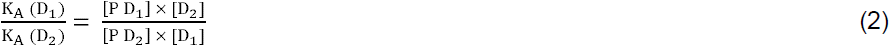

Relative binding affinities were determined based on three (wild-type SEP3_MI_ protein) or two (SEP3_MI_ R3A and SEP3_MI_ R3K proteins) different protein isolates, respectively. Measurements with each protein isolate were replicated three times. The DNA probes to be individually tested were labeled with the fluorescent dye Cy5. The chosen standard DNA probe was labeled with the fluorescent dye 6-Fam and contains the CArG-box sequence 5’-CCATTAATGG-3’. As a control, we used a reference labeled with the fluorescent dye DY-490 and containing the CArG-box sequence 5’-CCAAAAAAGG-3’. These control measurements were done with two different wild-type protein isolates and with one protein isolate for SEP3_MI_ R3A and SEP3_MI_ R3K, respectively, with only one replicate per sample. Highly similar results were obtained for both references (5’-CCATTAATGG-3’ labeled with 6-Fam and 5’- CCAAAAAAGG-3’ labeled with DY-490).

### Saturation binding assay

To determine the DNA-binding affinity of the SEP3_MI_ protein dimers, radioactively labeled DNA probes were used in electrophoretic mobility shift assays (EMSAs). In these assays, a constant amount of DNA was titrated against increasing concentrations of protein. Experiments were replicated four times.

Fractional occupancy of DNA-binding sites is defined as (adapted from (Smart and Hodgson, 2008)):

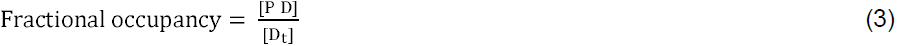

where [D_t_] is the total concentration of DNA-binding sites and [P D] the concentration of the protein-DNA complex. The total DNA concentration [D_t_] can be also phrased as:

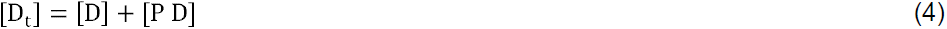

Combining equations (3) and (4) leads to:

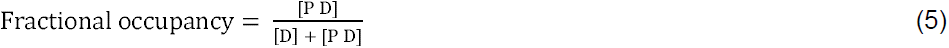

Equation (1) (here with K_A,dim_ – representing K_A_ for a protein dimer) can be rearranged as:

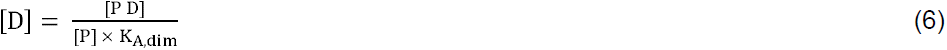

Combining equations (5) and (6) yields:

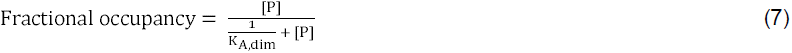

The concentration of free protein [P] is close to the total protein concentration [P_t_], when the DNA concentration is much lower than the inverse of the association constant (i.e. the dissociation constant):

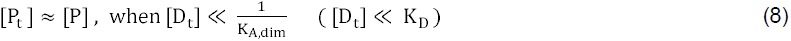

In a control experiment different DNA concentrations were tested to get a rough estimate of the K_D_ value. The DNA concentration finally used for the assay was 0.1 nM and thus much smaller than the K_D_ values which varied between 15 to 500 nM.

Substituting [P] by [P_t_] in Equation (7) yields:

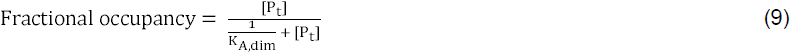

Plotting the fractional occupancy versus [P_t_] enables the determination of K_A,dim_ using a non-linear regression analysis with Equation (9). This was done in GraphPad Prism 6 (“Receptor binding – Saturation binding – Equation: One site – Specific binding”).

### Calculation of binding energies

The Gibbs energy ΔG for the binding of a SEP3_MI_ dimer to DNA was calculated using the equation:

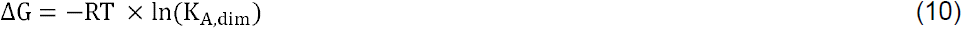

with R being the gas constant (8.314 · 10^−3^ kJ/(K·mol)) and T the absolute temperature (295.15 K; corresponding to 22 °C, the temperature at which the protein-DNA reaction was incubated). Apparent binding affinities K_A,dim_ to the probe with the CArG-box sequence 5’-CCATTAATGG-3’ were determined with the saturation binding assay. For the other probes relative binding constants were taken from the QuMFRA assay (Supplemental Data Set 2) and apparent binding affinities were calculated using the binding affinity K_A,dim_ of the CArG-box probe 5’-CCATTAATGG- 3’ as a reference value. Using Equation (11) binding energies were calculated based on the binding affinities K_A,dim_ (Supplemental Data Set 3).

### Prediction of binding energies based on an additivity model

Binding energies were predicted based on the assumption that each base pair of the CArG-box contributes independently to the overall binding energy. The model was calibrated using the experimentally determined affinity to the CArG-box sequence 5’- CCAAAAAAGG-3’.

CArG-box sequences like 5’-CCAAAAAAGG-3’ and 5’-CCTTTTTTGG-3’ are pairs of inverted sequences and represent identical double-stranded DNA molecules. However, slight differences in binding affinity to SEP3_MI_ were detected for these pairs (Figure 1), most likely because the sequences flanking the CArG-boxes on the 5’ and 3’ side were different from each other (Supplemental Figure 1) and thus created a directionality in SEP3_MI_ binding that influenced binding affinities. We accounted for this influence of the flanking sequences in the way that we calibrated the model in only one probe orientation. ‘Orientation’ refers here to the specified identity of the flanking sequences at the respective 5’- or 3’-side relative to the CArG-box. Otherwise we ignored the differences in the flanking regions for our model.

The Gibbs free energies of SEP3_MI_ binding to CArG-box sequences that deviated from 5’-CCAAAAAAGG-3’ in single substitutions were subsequently subtracted from the binding energy to 5’-CCAAAAAAGG-3’; i.e. sequences with one T instead of an A at each of the six different positions in the AT-stretch were used to calculate “penalties” for the A-to-T substitution per position (Supplemental Data Set 3), as previously described (Stormo and Fields, 1998; Maerkl and Quake, 2007). By summing up the penalties for specific positions and subtracting those from the binding energy of the reference sequence 5’-CCAAAAAAGG-3’ the Gibbs free energies for SEP3_MI_ binding to other CArG-box sequences could be predicted (Supplemental Data Set 3). We assumed that penalties relative to the reference sequence should be kept to a minimum in the model calculations in order to reduce cumulative errors, e.g. due to orientation of the CArG-box. Thus, penalties for each pair of inverted CArG-box sequences were calculated and the lower penalty (higher binding energy) was used for both orientations.

For model evaluation and comparison the root mean square error of prediction (RMSEP) was calculated. The RMSEP is defined as:

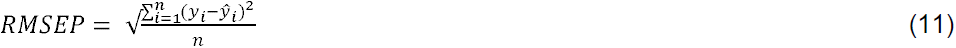

where y_i_ is the measured value of y (measured binding energy) for object i, ŷ_i_ is the y-value for object i predicted by the model under evaluation (predicted binding energy) and n is the number of objects for which pairs of yi and ŷ_i_ are available (number of measurements/ predictions).

### Analysis of DNA shape features

Structural features of the QuMFRA DNA probes (25 sequences) were predicted using the online tool DNAshape (Zhou et al., 2013). The following DNA shape parameters for dsDNA were predicted: the minor groove width (MGW), the roll angle (Roll), the propeller twist (ProT) and the helical twist (HelT). Either the parameters for the central base (in the case of MGW and ProT) or the central base pair steps (for Roll and HelT) were predicted based on a sliding-pentamer window.

For each CArG-box the shape parameters of the six central A/T bases (six parameters for MGW, six for ProT, seven for Roll and seven for HelT) were extracted (Supplemental Data Set 4).

### Analysis of ChIP-seq data

ChIP-seq data sets for SEP3 (Kaufmann et al., 2009), AP1 (Kaufmann et al., 2010) and FLC (Deng et al., 2011) were used here based on a reanalyzed form (Muiño et al., 2014) where for all three proteins ChIP-seq scores per CArG-box of the type 5’- CC(A/T)_6_GG-3’ for the complete *Arabidopsis thaliana* genome were given. We simplified these data sets by calculating for each protein mean ChIP-seq scores for the 64 different CArG-box sequences. Then we reduced the data set further by combining the pairs of inverted CArG-box sequences. We thus ignored possible orientation or flanking sequence effects on binding affinity. We used the mean value of these CArG-box pairs for our analysis, thereby reducing the 64 to 36 different CArG-box sequences. Finally, we calculated Spearman rank correlation values in order to elucidate whether higher values for binding affinities obtained by our gel shift assays as well as higher predicted binding energies obtained by the additivity model matched higher mean ChIP-seq scores.

### Accession numbers

Accession numbers are listed in Supplemental Table 1.

## Supplemental Data

**Supplemental Figure 1.** Domain structure of SEPALLATA3 (SEP3) and recombinant protein expression of SEP3_MI_.

**Supplemental Figure 2**. Lysis of *E. coli* cells and first purification step of SEP3_MI_ using a Ni sepharose column.

**Supplemental Figure 3**. Second purification step of SEP3_MI_ using a heparin sepharose column.

**Supplemental Figure 4**. Third purification step of SEP3_MI_ using a size exclusion chromatography column.

**Supplemental Figure 5**. Quantitative Multiple Fluorescence Relative Affinity (QuMFRA) assay results.

**Supplemental Figure 6.** The preference for A-tract sequences is not dependent on the flanking sequences.

**Supplemental Figure 7**. Saturation-binding assay.

**Supplemental Table 1**. DNA and protein sequences of SEP3_MI_, SEP3_MI_ R3A and SEP3_MI_ R3K.

**Supplemental Data Set 1**. DNA probe design for gel shift assays.

**Supplemental Data Set 2**. QuMFRA assay results.

**Supplemental Data Set 3**. Predictions from the additivity model.

**Supplemental Data Set 4**. DNA structural parameters predicted by DNAshape

**Supplemental Data Set 5.** Comparison of *in vitro* QuMFRA data with *in vivo* ChIP-seq data.

## ACKNOWLEDGEMENTS

We thank Jens Schumacher for help with statistical analyses. This work was supported by a fellowship from the International Leibniz Research School for Microbial and Biomolecular Interactions (ILRS Jena) which is part of the Jena School for Microbial Communication (JSMC) [to S.G.]; a fellowship from the Carl-Zeiss-Stiftung [to R.M.] and the Deutsche Forschungsgemeinschaft [grant number TH417/5-3] to [G.T and R.M.].

### AUTHOR CONTRIBUTIONS

R.M. and G.T. designed the research; S.G. and C.G. performed research; R.M., S.G. and F.R. analyzed data; S.G. and R.M. wrote the manuscript.

